# Evidence of Presence and Activation of the two Cell Cycle Kinases Nek6 and Nek7 in the ciliated Neuroretina

**DOI:** 10.1101/2020.03.27.012724

**Authors:** Kerstin M Janisch, J Mie Kasanuki, Richard J Davis, Stephen H Tsang

**Affiliations:** Bernard & Shirlee Brown Glaucoma Laboratory, Harkness Eye Institute, Department of Pathology, College of Physicians and Surgeons Columbia University Medical Center, New York, NY 10032, USA

**Keywords:** NEK kinase, retina, GPCR signalling, photoreceptor, cilia

## Abstract

The serine/threonine NIMA kinases are widely found in eukaryotes. They are cell-cycle kinases that are associated with centrosomes and spindle apparatus and cilia. In cilia, NIMA kinases are reported to play a role in cilia length maintenance and deflagelation. Here we focus on the two Nek homologs, Nek6 and Nek7, and their potential role in retina. We report for the first-time expression of *nek6* and *nek7* mRNA and protein in retinal tissue. In particular, we detect localisation of these kinases to photoreceptors outer segments. Moreover, we are able to show a light-dependent phosphorylation of the activation loop (serine 206) of Nek6/7 in rod outer segments, suggesting activation of these kinases is downstream of the phototransduction pathway. Indeed, we demonstrate that Nek6/7 phosphorylation in the retina is dependent on *Grk1* function. Furthermore, Nek6/7 phosphorylation can be stimulated in the brain by opiate drugs, suggesting that activation of Nek6/7 lies downstream of G protein coupled receptors activation, in general. Nek6/7 may couple photoreception with outer segment biogenesis through phosphorylation of downstream substrates, which may affect the microtubules of the axoneme.

## Introduction

Kinases are important regulators in cellular metabolism. Phosphorylation, as one of the main post-translational modifications of proteins, plays a crucial part in regulatory processes. Phosphorylation and dephosphorylation act as on/off switches for a variety of enzymes and receptors (Helmreich, 2001). The lack of a kinase or its up-regulation can have multiple implications ranging from developmental impairments and inflammatory diseases to cancer (http://www.cellsignal.com/reference/kinase_disease.html).

The NIMA (Never in Mitosis A)-related protein kinases (Nek) are serine/threonine kinases associated with the cell cycle (Malumbres and Barbacid, 2007; O’Regan et al., 2007; Quarmby and Mahjoub, 2005). The Nek2 clade is associated with G2/M regulation and centrosome separation (Li & Li, 2006); whereas, Nek1 and Nek8 contribute to ciliary function (Mahjoub et al., 2005; Otto et al., 2008). The clade of Nek6 and Nek7 has been linked to mitotic progression, spindle assembly and microtubule nucleation (Kim et al., 2007; Quarmby and Mahjoub, 2005; Yin et al., 2003; Yissachar et al., 2006). Nek6 and Nek7 are highly homologous; they share 87% amino acid identity (Belham et al., 2003; Quarmby and Mahjoub, 2005). The kinases are activated by phosphorylation at serine206 (S206), which is found within the activation loop (amino acids 183-192 and 197-210) of both kinases. The suggested upstream kinase for activating Nek6/7 is Nek9/Nercc1 (Belham et al., 2003; O’Regan et al., 2007), but other upstream kinases have been published such as Chk1/2 or ATM (Lee, et al., 2008; Moraes, et al., 2015; Tan et al., 2017). In studies with recombinant, as well as, exogenously NIMA kinase expressing cells, the control mechanisms of Nek6 and Nek7 differ under physiological conditions (Minoguchi et al., 2003). Additionally, the distinctive expression patterns in various tissues suggests different properties for these kinases *in vivo* (Feige and Motro, 2002). Interestingly, the clade of Nek6/7 is not found fungi, suggesting a role of Nek6/7 in ciliary function (O’Regan et al., 2007; Quarmby and Mahjoub, 2005). Nek7 has recently been linked within the inflammasome complex involving NLRP3 (He et al., 2016), which plays a role in bystander cone photoreceptor cell death (Viringipurampeer et al., 2018).

The rod photoreceptor is a specialized non-motile cilium. The outer segment contains the machinery for light detection and is connected with a connecting cilia to the inner segment, which holds all organelles for biosynthesis, (Roepman and Wolfrum, 2007). The basal body of cilia derives originally from the mother centrosomes of a cell (Greiner et al., 1981). We, therefore, hypothesized that NIMA kinases, connected with centrosomes and spindle apparatus can also be found in the photoreceptor. To investigate their presence and possible activation, we took a system biology approach using molecular biology and biochemistry.

## Results

### RT-PCR

NIMA kinases have been reported to be involved with ciliogenesis. Several eye diseases are now classified as ciliopathies. To investigate transcription of these kinases in mouse retinae, we performed RT-PCR and qRT-PCR on mouse retinae and olfactory bulbs as control tissue for the transcription of *nek6*, *nek7*, *nek8* and *nek9*. Total mRNA of all four kinases is found (**Figure 1**) in murine retina, as well as in murine olfactory bulbs. *nek6* and *nek7* transcription in the olfactory bulbs has been previously described (Feige and Motro, 2002) and were chosen as positive controls. Lane 3 in each set is the negative control with no cDNA template added. qRT-PCR was performed to analyse the level of RNA transcribed in both murine olfactory bulbs and retinae. The housekeeping gene *β-actin* was used in both tissues to standardize RNA levels (Dheda et al., 2004). **Table 1** shows the qRT-PCR data for olfactory bulbs and retina in the 2^−ΔCt^ format (Schmittgen and Livak, 2008). Olfactory bulbs had higher levels of *nek6* and *nek7* than mouse retina. The only difference was that *nek7* was highly statistically significant (p < 0.01). The difference of the *nek6* levels was 6.9-fold; whereas, the *nek7* levels differed 4.5-fold (**Table 2**). *nek8* was not detectable in olfactory bulb tissue; *nek9* RNA levels were 4.4-fold higher in the retinal tissue in comparison to olfactory bulb tissue, a difference which was not statistically significant (**Table 1 and 2**). In the mouse retina, the RNA levels of *nek6* and *nek7* differed 130.9-fold, which is very highly significant (p < 0.001; **Table 3**). Even though there was a 6.8-fold difference between *nek6* and *nek8* levels, there is no statistic significance (**Table 3**). *nek7* RNA levels were 1.3-fold higher than *nek9* at a significant level (p < 0.05; **Table 3**). In olfactory bulbs, *nek6* and *nek9* levels were very highly significant different from *nek7* levels, with 54.5-fold and 25.4-fold difference, respectively (**Table 4**). RNA levels for *nek6* were 2.2-fold lower than *nek9* levels, a highly significant difference (p < 0.001; **Table 4**).

**Table 1.** qRT-PCR results for mouse olfactory bulbs and retina presented as Avg 2_−ΔCt_ (n = 6) with stdev for RNA levels of *nek6*, *nek7*, *nek8* and *nek9*. n.d., not detectable

|  | OLFACTORY BULBS |  | RETINA |  |
| --- | --- | --- | --- | --- |
| | Avg 2-ΔCt | Stdev | Avg 2-ΔCt | Stdev |
| *nek6* | 3.0 x 10<sup>-3</sup> | 4.4 x 10<sup>-4</sup> | 4.4 x 10<sup>-4</sup> | 3.5 x 10<sup>-4</sup> |
| *nek7* | 2.6 x 10<sup>-1</sup> | 1.8 x 10<sup>-2</sup> | 5.7 x 10<sup>-2</sup> | 4.7 x 10<sup>-3</sup> |
| *nek8* | n.d. | n.d. | 2.9 x 10<sup>-3</sup> | 2.7 x 10<sup>-4</sup> |
| *nek9* | 1.0 x 10<sup>-2</sup> | 2.4 x 10<sup>-3</sup> | 4.4 x 10<sup>-2</sup> | 1.4 x 10<sup>-3</sup> |

**Table 2.** Statistical calculations for RNA levels of different *nek* transcripts in murine retinal tissue compared to olfactory bulbs. Probability was calculated using F-test. Significance given as: *, significantly different (p < 0.05); **, highly significantly different (p < 0.01); ***, very highly significantly different (p < 0.001); # since *nek8* is undetectable, the value for lim_OLFACTORYBULBSvsRETINA_(2.9 × 10_−3_/0) ⇒∞

| OLFACTORY BULBS vs. RETINA |  |  |  |
| --- | --- | --- | --- |
|  | F-test probability | Significance | Fold difference |
| *nek6* | 0.636 |  | 6.9 |
| *nek7* | 0.0097 | ** | 4.5 |
| *nek8* | | *** | ∞ # |
| *nek9* | 0.268 |  | 4.4 |

**Table 3.** Statistical calculations for RNA levels of different *nek* transcripts in murine retinal tissue. Probability was calculated using F-test. Significance given as: *, significantly different (p < 0.05); **, highly significantly different (p < 0.01); ***, very highly significantly different (p < 0.001)

|  | F-test probability | Significance | Fold difference |
| --- | --- | --- | --- |
| *nek6* vs. *nek7* | 0.00003 | *** | 130.9 |
| *nek6* vs. *nek8* | 0.589 |  | 6.8 |
| *nek6* vs. *nek9* | 0.009 | ** | 100.8 |
| *nek7* vs. *nek8* | 0.00001 | *** | 19.3 |
| *nek7* vs. *nek9* | 0.019 | * | 1.3 |
| *nek8* vs. *nek9* | 0.003 | ** | 14.9 |

**Table 4.** Statistical calculations for RNA levels of different *nek* transcripts in murine olfactory bulbs. Probability was calculated using F-test. Significance given as: *, significantly different (p < 0.05); **, highly significantly different (p < 0.01); ***, very highly significantly different (p < 0.001); # since *nek8* is un-detectable, the values for lim_*nek6*vs*nek8*_(3.0 × 10_−3_/0), lim_*nek7*vs*nek8*_(2.6 × 10_−1_/0) and lim_*nek9*vs*nek8*_(1.0 × 10_−2_/0) ⇒∞

|  | F-test probability | Significance | Fold difference |
| --- | --- | --- | --- |
| *nek6* vs. *nek7* | 0.0000001 | *** | 54.5 |
| *nek6* vs. *nek8* | | *** | ∞ |
| *nek6* vs. *nek9* | 0.002 | ** | 2.2 |
| *nek7* vs. *nek8* | | *** | ∞ |
| *nek7* vs. *nek9* | 0.00039 | *** | 25.4 |
| *nek8* vs. *nek9* | | *** | ∞ |

**Figure 1.**
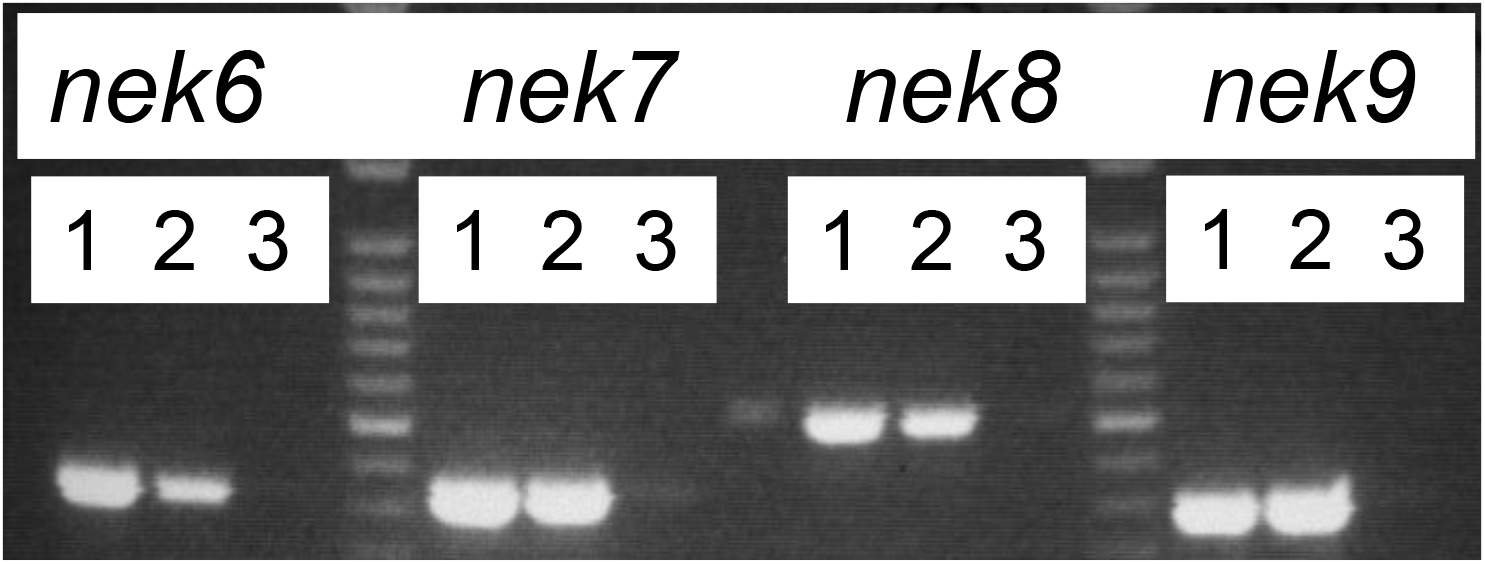
Presence of NIMA kinase mRNA in retina and olfactory bulbs. RT-PCR for *nek6*, *nek7*, *nek8* and *nek9* kinases in mouse olfactory bulbs and retinae. 1, olfactory bulbs; 2, retina; 3, water

### 2D-Immunoblots

The high RNA levels of Nek7 in the retina with qRT-PCR led us to investigate the expression of the two NIMA kinases Nek6 and 7, as they are highly homologous. To separate both kinases and visualize possible post-translational modifications we employed 2D-electrophoreses. Rhodopsin, the most abundant protein in photoreceptor, was used as an internal control and to verify no possible cross-reaction. As **Figure 2** shows, Nek6 and Nek7 were detected in dark- and light-adapted cow rod outersegment preparations. Probing with anti-Nek7 antibody resulted in two spots at around 36 kDa (**Figure 2A** and **D**). The spot shifting to the more acidic pH suggested a post-translational modification of the protein, and the intensity of this spot was increased in the light-adapted sample (**Figure 2A** and **D**). After stripping, re-probing with anti-Nek6 antibody revealed two spots at similar locations. Again, the shift of one spot to the acidic site suggested post-translational modifications. When probed with the Nek6 antibody, there appeared to be no difference in spot intensity between the dark- and light-adapted samples (**Figure 2B** and **E**). Re-probing of the membranes with a rhodopsin antibody showed a different pattern for this protein. The intensity of the protein increased in light-adapted samples. Rhodopsin is a highly post-translational modified protein; thus, no concrete spot formation was observable. It appeared as a band on the 2D-blot (**Figure 2C** and **F**). All three images of Nek6, Nek7 and rhodopsin were superimposed for the dark- and light-adapted samples (**Figure 3**). The two NIMA kinases and rhodopsin could clearly be distinguished. Furthermore, merged images showed Nek6 and Nek7 overlaying each other, which complies with their high homology (**Figure 3**, dark blue, Nek6; light blue, Nek7).

**Figure 2:**
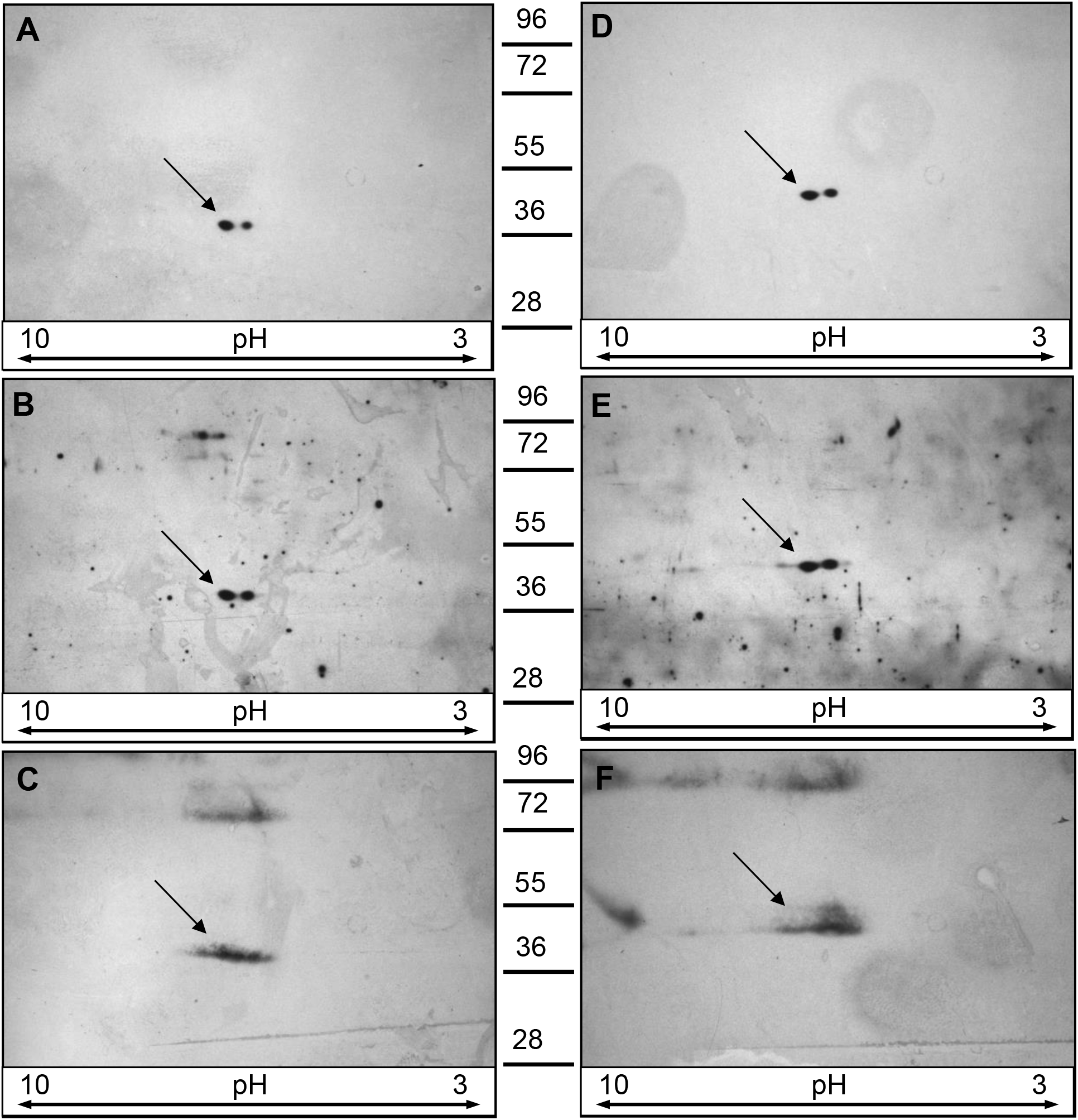
2D-immunoblots on dark- and light-adapted cow rod outersegments. Membranes shown were probed with antibodies against Nek6, Nek7 and rhodopsin. Proteins are indicated with an arrow. A, D anti-Nek7; B, E, anti-Nek6; C, F anti-rhodopsin; A, B, C, dark-adapted sample; D, E, F, light-adapted sample; 50 μg total protein loaded

**Figure 3.**
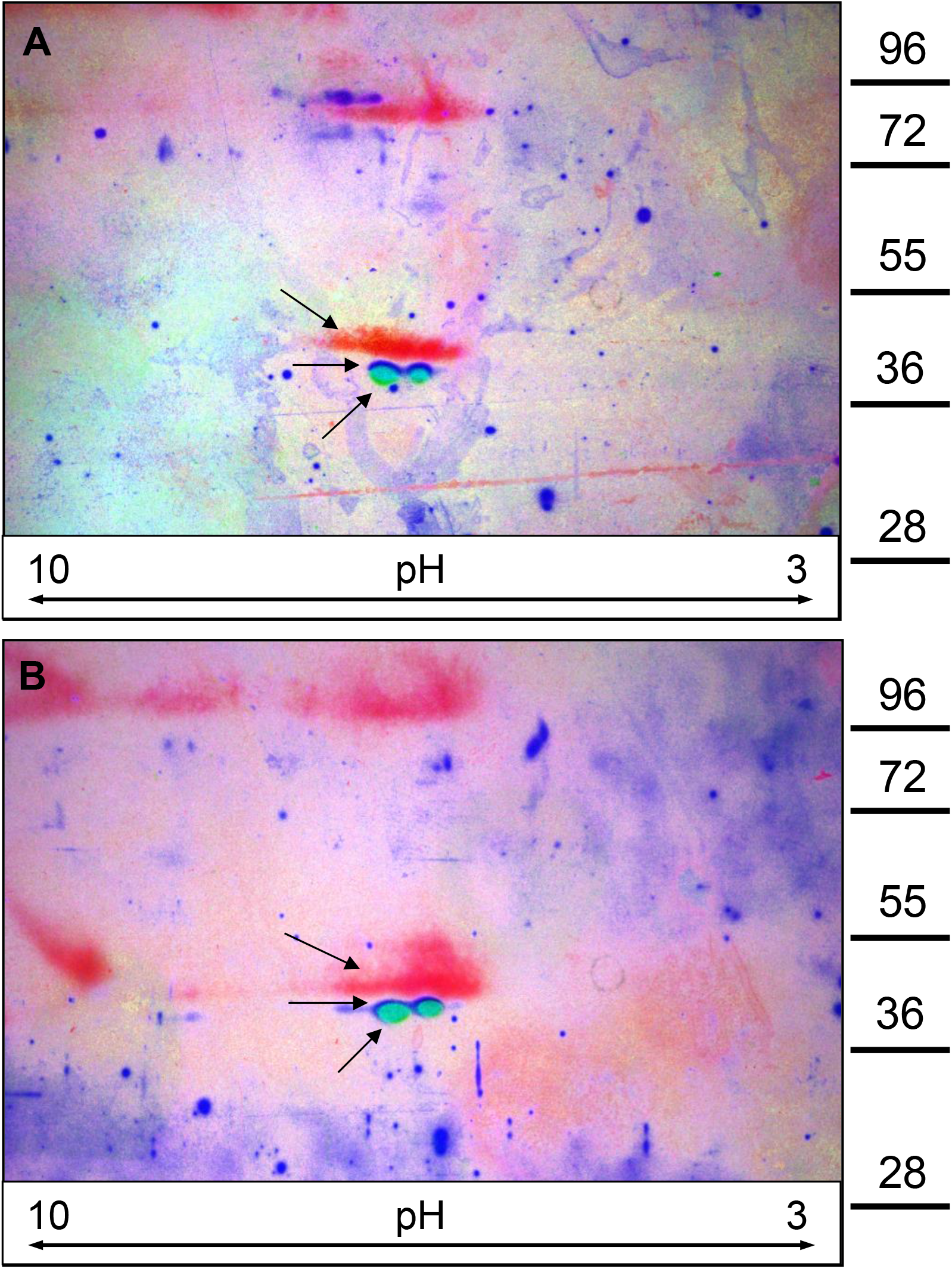
Overlay of 2D-immunoblots of dark- and light-adapted cow rod outersegments. A, dark-adapted; B, light-adapted; red, rhodopsin; blue, anti-Nek6; light blue, anti-Nek7; arrows indicate proteins, 50 μg total protein loaded

### Immunohistochemistry

The 2D-immunoblot findings encouraged us to investigate further the distribution of the two kinases Nek6 and Nek7 in dark- and light-adapted retina. We probed frozen sections of dark- and light-adapted retinae (on the same slide) with antibodies against Nek6, Nek7, and phosphoNek6/7. The exposure times between the dark- and light-adapted samples were kept the same for each antibody.

As seen **in Figure 4A**, Nek6 exhibits diffuse staining in the OS, IS, OPL and IPL in dark-adapted retinae, which increases in signal in the light-adapted retina (**Figure 4D**). Nek7 has a slightly higher expression signal in the dark-adapted retina (**Figure 4B**) than Nek6. It is also clearly expressed in IPL, OPL and IS in both dark- and light-adapted retinae (**Figure 4B** and **E**) but is barely detectable in the the OS of the dark-adapted retina (**Figure 4B**). In the light-adapted sections (**Figure 4E**), there is a clear increase in Nek7 signal, which would respond to an increase in protein levels.

**Figure 4.**
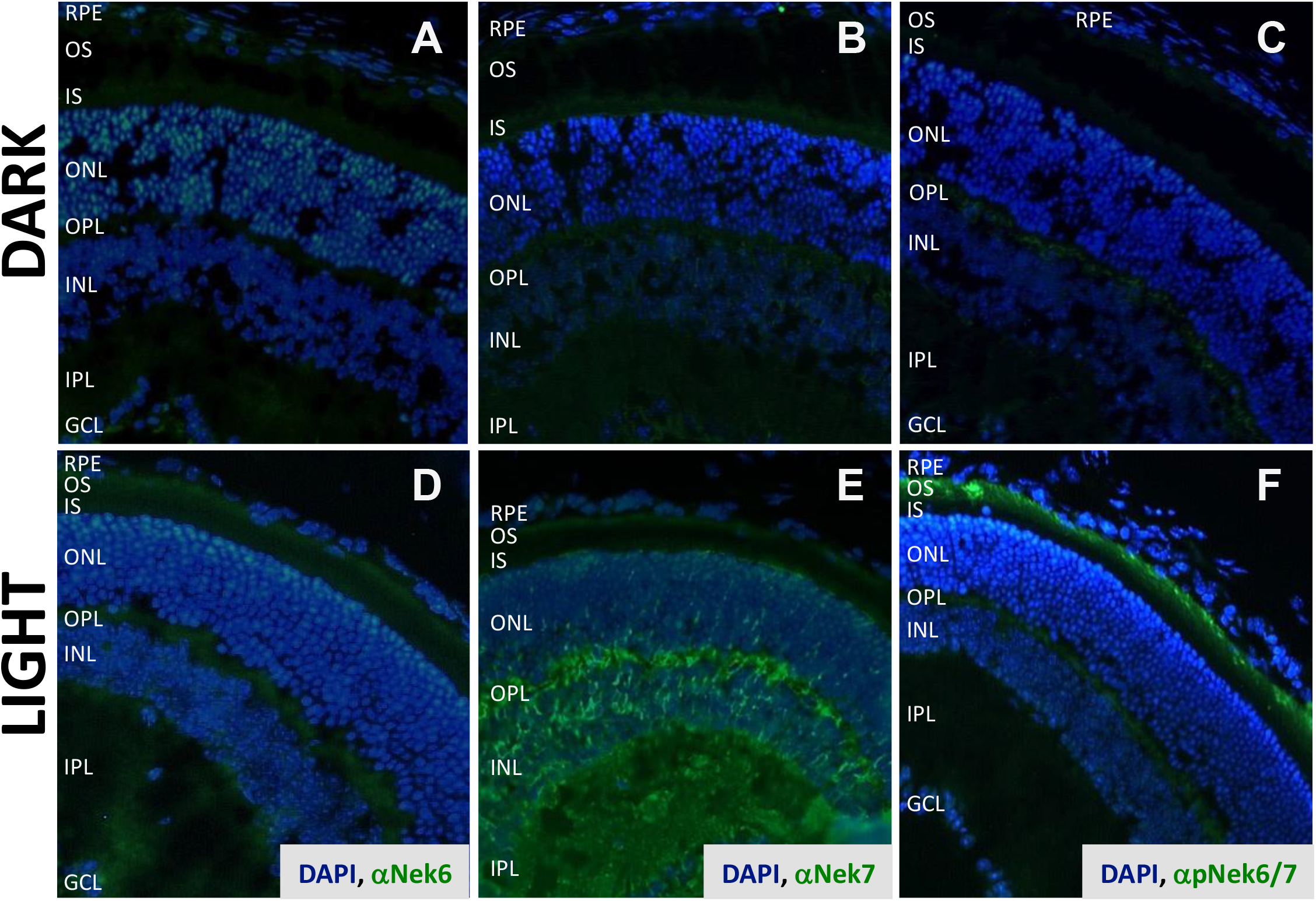
Immunohistochemistry (IHC) of frozen sections of dark- and light-adapted mouse retinae. dark- and light-adapted sections were on the same slide; A: dark-adapted retina with anti-Nek6 + DAPI; B: dark-adapted retina with anti-Nek7 + DAPI; C: dark-adapted retina with anti-posphoNek6/7+ DAPI; D: light-adapted retina with anti-Nek6 + DAPI; E: light-adapted retina with anti-Nek7 + DAPI; F: light-adapted retina with anti-phosphoNek6/7 + DAPI; RPE, retinal pigment epithelium; OS, outer segment; IS, inner segment; ONL, outer nuclear layer; OPL, outer plexiform layer; INL, inner nuclear layer; IPL, inner plexiform layer; GCL, ganglion cell layer

Thanks to the generous gift of a phospho-Nek6/7 antibody from Dr Roig Amorós, we were able to investigate the observed post-translational modification seen in the 2D-immunoblot. The phospho-Nek antibody was raised against a peptide representing amino acids 203–213 including phospho-serine206 (pS206) of the active core (amino acid sequence: AAHpSLVGTPYY; Dr Joan Roig Amorós, personal communication). Due to the high homology of Nek6 and Nek7 (Belham et al., 2003; Minoguchi et al., 2003; O’Regan et al., 2007; Quarmby and Mahjoub, 2005), this sequence occurs in both kinases and the detected signal of the antibody can be attributed to either of the activated kinases.

The dark-adapted retina exhibits some signal within the OPL, when probed with the pNek6/7 antibody (**Figure 4C**). Whereas in the light-adapted retina the phosphorylation could clearly be observed as a strong signal in the photoreceptors (**Figure 4F**). A double-staining with centrin-1 (data not shown) verified, that the phosphorylation of Nek6/7 occurred within the photoreceptors in the light-adapted sample.

### Immunoblots

The IHC findings led us to investigate the phosphorylation of Nek6/7 further. Retinal extracts of MF1 mice that were subjected to different light exposures were analysed with the phosphoNek6/7 antibody. The immunoblot analysis confirmed the absence of the phosphorylation in dark-adapted samples, with the onset of the phosphorylation occurring within 1-minute post-light exposure. The phosphorylation increased over time and reached a maximum level when the retina was completely light-adapted (= bleached) (**Figure 5A**). The membranes were then stripped and re-probed with β-actin to ensure constant loading of the samples **(Figure 5B**).

**Figure 5.**
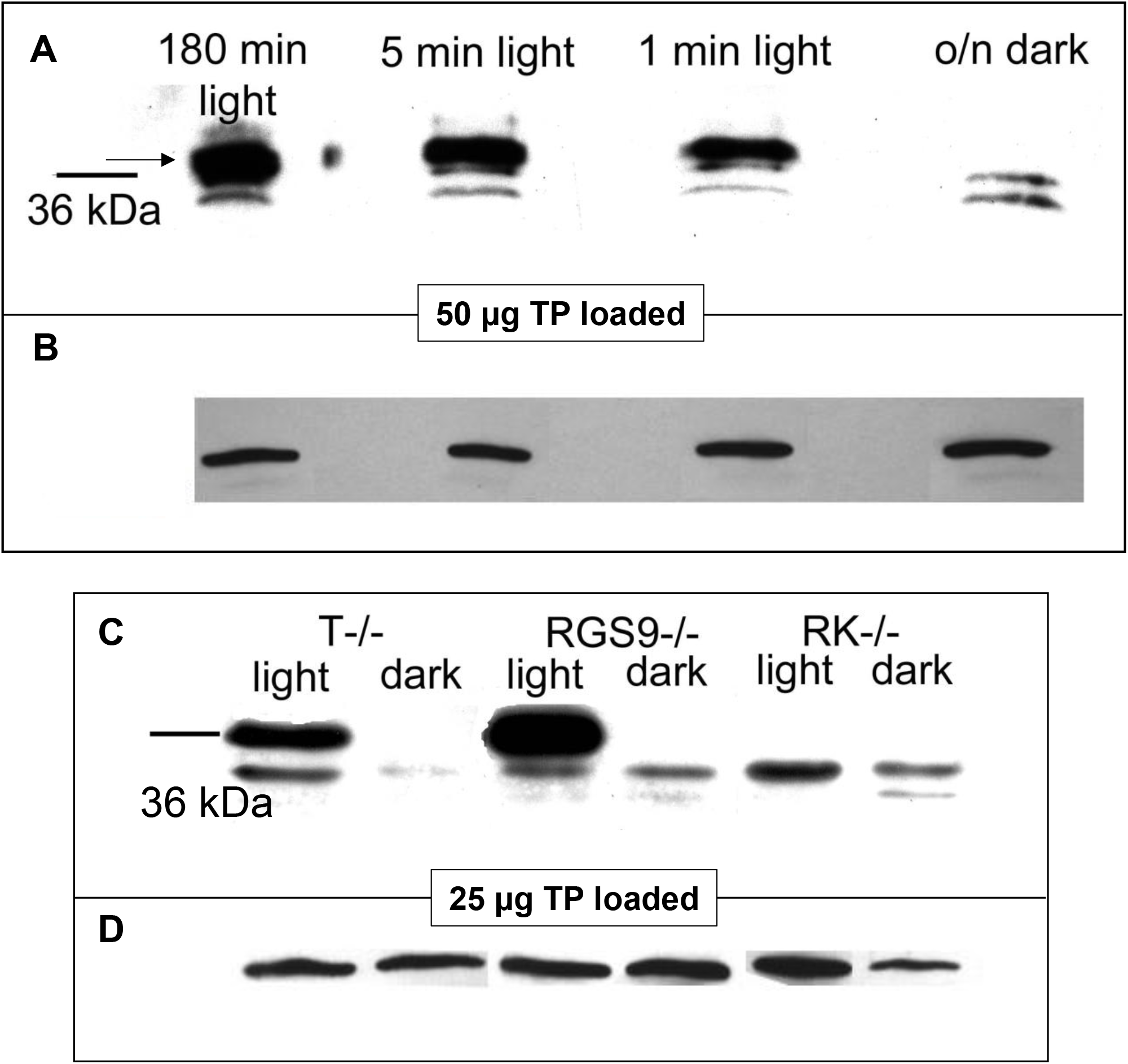
Immunoblots of retinal extracts. TP, total protein; A, extracts from retina with different light exposure times; B, β-actin loading control; C, dark- and light-adapted retinal extracts of different knock-out mice retinae; D, β-actin loading control; T−/−, transducin knock-out mouse; RGS9−/−, regulator of G-protein signalling 9 knock-out mouse; RK−/−, rhodopsin kinase knock-out mouse; arrow indicates phospho-protein

This result encouraged us to investigate mouse lines with either transducin knockouts (T−/−), regulator of G-protein signalling 9 knockouts (RGS9−/−) and rhodopsin kinase knockouts (RK −/−). Dark- and light-adapted retinae of knockouts were a generous gift of Jason Chen. The phosphorylation of Nek6/7 was not affected in T−/− mice; whereas, RGS9−/− mice showed an increased phosphorylation in the light. Interestingly, RK−/− mice exhibited no phosphorylation in dark or light (**Figure 5C**). **Figure 5D** shows the β-actin reprobing as a loading control for this sample set.

With these results, we wanted to investigate, whether the phosphorylation of Nek6/7 occurs in other G-protein coupled signalling cascades. For this, we intraperitoneally injected mice with buprenorphine, cocaine, amphetamine, norepinephrine or saline (as control) to stimulate opioid, dopaminergic and adrenergic receptors in various regions of the brain (striatum, substantia nigra). **Figure 6** shows the densitometry results of the immunoblots done on these samples. With buprenorphine injections there was no significant change in Nek6/7 phosphorylation in any of the aforementioned brain regions. Cocaine and amphetamine exhibited similar results with Nek6/7 phosphorylation. In the striatum, both drugs increased the Nek6/7 phosphorylation; this was also the case in the substantia nigra. Norepinephrine injections increased the phosphorylation in the striatum as seen with cocaine and amphetamine. In contrast, norepinephrine decreased the phosphorylation of Nek6/7 in the substantia nigra (**Figure 6**).

**Figure 6.**
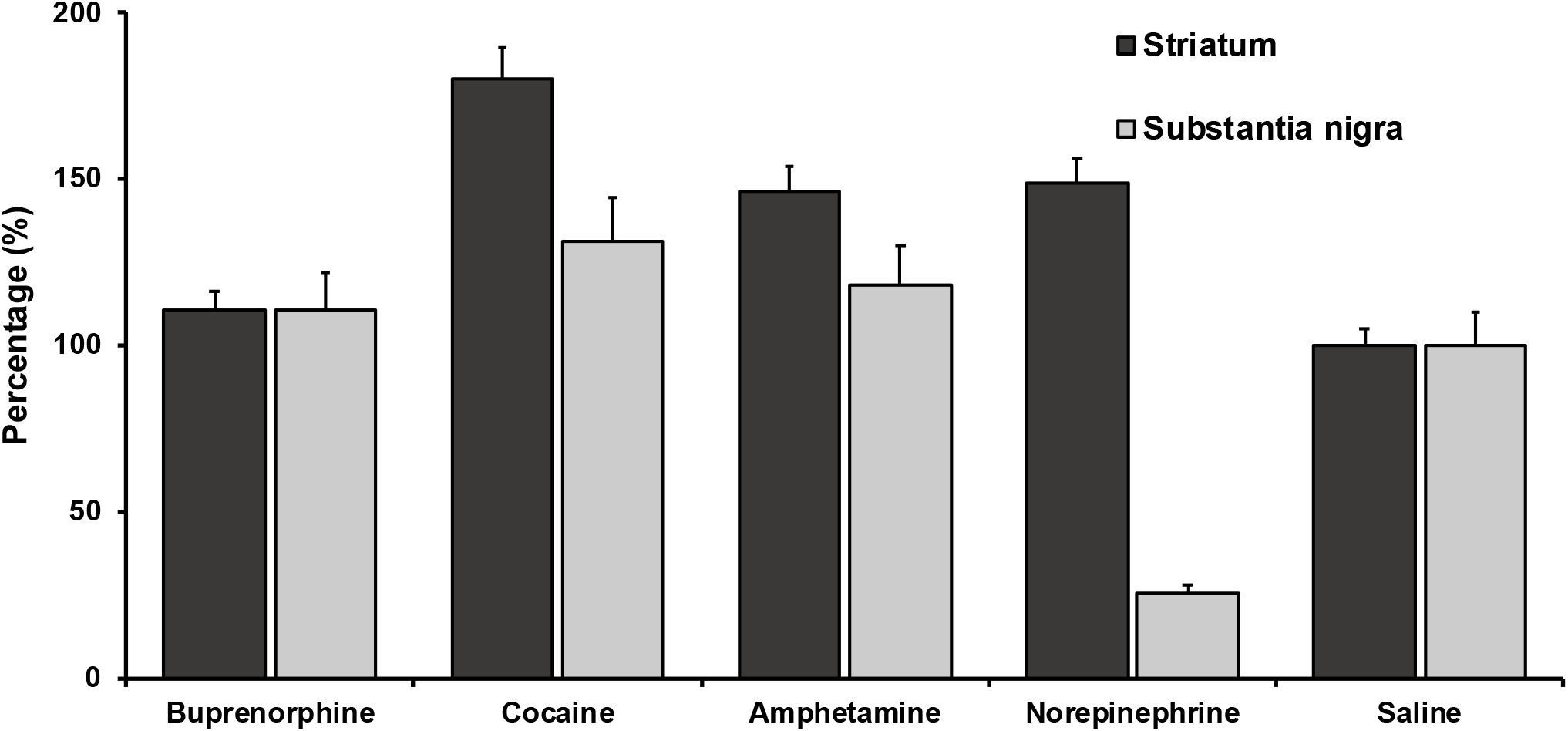
Densitometry results of Nek6/7 phosphorylation in striatum and substantia nigra samples of injected mice

## Discussion

NIMA kinases are cell cycle kinases that have been associated with several ciliopathies and cancers (Afzelius, 2004; de Carcer et al., 2007; Quarmby and Mahjoub, 2005). The kinases Nek1 and Nek8, for example, are linked with polycystic kidney disease and nephronophthisis, two ciliopathies of renal cells (Mahjoub et al., 2005; Otto et al., 2008). Photoreceptor cells are, like kidney cells, primary cilia (Calvet, 2003). Our RT- and qRT-PCR results reveal for the first time the presence of *nek6*, *nek7, nek8* and *nek9* mRNA in mouse retina. *nek7* is predominant followed by *nek9*, which transcribes the proposed upstream kinase (Belham et al., 2003). *nek6* and *nek8* are found at very low levels. The mRNA distribution in the control tissue, the olfactory bulbs, is consistent with the already published distribution (Feige and Motro, 2002).

Since NIMA kinases are expressed on a low level in tissue in general (Lu and Hunter, 1995), we chose cow rod preparations for higher yields. The expression patterns of Nek6 and Nek7 of dark- and light-adapted cow rod outer segments illustrates the phosphorylation of the kinases. An addition of a phosphate group commonly shifts a protein spot towards the acidic pH (Link, 1999). The phosphorylation of Nek7 is more prominent and follows a dark/light switch; whereas, this post-translational modification only slightly changes with a dark/light switch in Nek6. Based on spot size, Nek7 kinase levels are slightly higher than Nek6. Rhodopsin that is clearly distinguishable from the kinases has a streak like pattern.

Both Nek 6 and Nek7 are diffusely distributed throughout the retina. There is a clear accumulation in the inner plexiform (IPL), outer plexiform (OPL) and inner segment (IS) layers in the dark- and light-adapted retina. The signals in these layers increase in intensity when probed for the whole protein in the light-adapted sample. The phosphorylated form of the kinase can be found in the OPL in the dark, and in the OPL and outer segment (OS) layer in the light. The observed signal of phosphorylation in the OS is clearly light dependent.

The “on-switch” of the kinase occurring within one minute of light exposure is a proof of the quickness of phosphorylation as a response to stimuli. In regulatory processes, signals need to be relayed quickly and efficiently. The use of phosphorylation as a main switch mechanism in signal transduction is well known (Helmreich, 2001; Walsh, 2008).

The lack of Nek6/7 phosphorylation in RK−/− mice prompted the investigation of a possible activation of Nek6/7 occurring in other GPCR signalling cascades. The brain provides several rhodopsin-like GPCR signalling cascades in various regions that have published expression of both kinases (Feige and Motro, 2002).

The clade of Nek kinases is not only associated with cell cycle control and spindle formation, but also with ciliary function (Kim et al., 2007; Mahjoub et al., 2005; O’Regan and Fry, 2009; Otto et al., 2008; Quarmby and Mahjoub, 2005; Wloga, 2005; Yin et al., 2003; Yissachar et al., 2006). Photoreceptors are specialised primary cilia and retinal neurons are ciliated (Adams et al., 2007). Both striatum and substantia nigra belong to the basal ganglia (Handel et al., 1999; Schulz et al., 2000; Siegel et al., 1998). Neurons of the basal ganglia have been reported to be ciliated (Handel et al., 1999; Schulz et al., 2000). Several groups reported (Berbari et al., 2009; Handel et al., 1999; Schulz et al., 2000) that neurotransmitter receptors (somatostatin and serotonin 5-HT_6_) are specifically confined in neural cilia in brain. They argue that there might be a function for the cilia in the neural signalling (Berbari et al., 2009; Pan et al., 2005; Veland et al., 2009).

Norepinephrine is a ligand for α_1_, α_2_ and β_1_ adrenergic receptor, which belongs to the rhodopsin-like GPCR subfamily A17 (EMBL# IPR 002233). The phosphorylation, and therefore Nek activation, downstream the adrenergic receptor could only be seen in the striatum. Buprenorphine stimulates the μ-opiod receptor belonging to the rhodopsin-like GPCR subfamily A4 (EMBL# IPR 001418). In none of the chosen brain regions was any Nek activation detectable with buprenorphine injections. The drugs amphetamine and cocaine exhibit their action on the dopamine receptors D1 and D2 through the inhibition of the dopamine active transporter. The dopamine receptors are categorized as GPCR subfamily A16 receptors (EMBL# IPR 000929). Nek6/7 are activated in the striatum and substantia nigra, when the dopamine receptors are stimulated (Siegel et al., 1998).

Recently, Nek7 has been found to be essential for MT nucleation (Kim et al., 2007), MT stabilization through Eg5 (Freixo et al., 2018), and to play a role in MT stability and dynamics (Cohen et al., 2013). The authors showed that Nek7 increases the growth and depolymerization velocity of MT and lengthens time of catastrophe. Several other NIMA kinases have been found to play a role in deflagelation and cilia length maintenance (Gaertig and Wloga, 2008). We hypothesize that these NIMA kinases in the photoreceptor cilia help maintain proper photoreceptor length by keeping the axoneme of the OS dynamic. The shedding of the tip of the photoreceptor follows a circadian rhythm linked to light onset (Hollyfield and Gasinger, 1978). A direct linkage between the external signal and stimuli of the GPCR signalling cascade would help maintain consistent cell length.

The results presented here led us to formulate the following model for a signaling cascade for Nek6/7 kinases (**Figure 7**). The main activation of Nek is through GRK1; nevertheless, there could be a percentage of NIMA kinase activation in a GPCR-independent matter (Gurevich & Gurevich, 2019). This would be supported by the results in **Figure 5C**, where transducin knockout (T−/−) retinae still show a phosphorylation of the kinase. At least in photoreceptors, an activation through oxidative damage needs to be considered and further investigated, as it has been shown that Nek7 exhibits protection of oxidative damage in telomeres (Tan et al., 2017).

**Figure 7:**
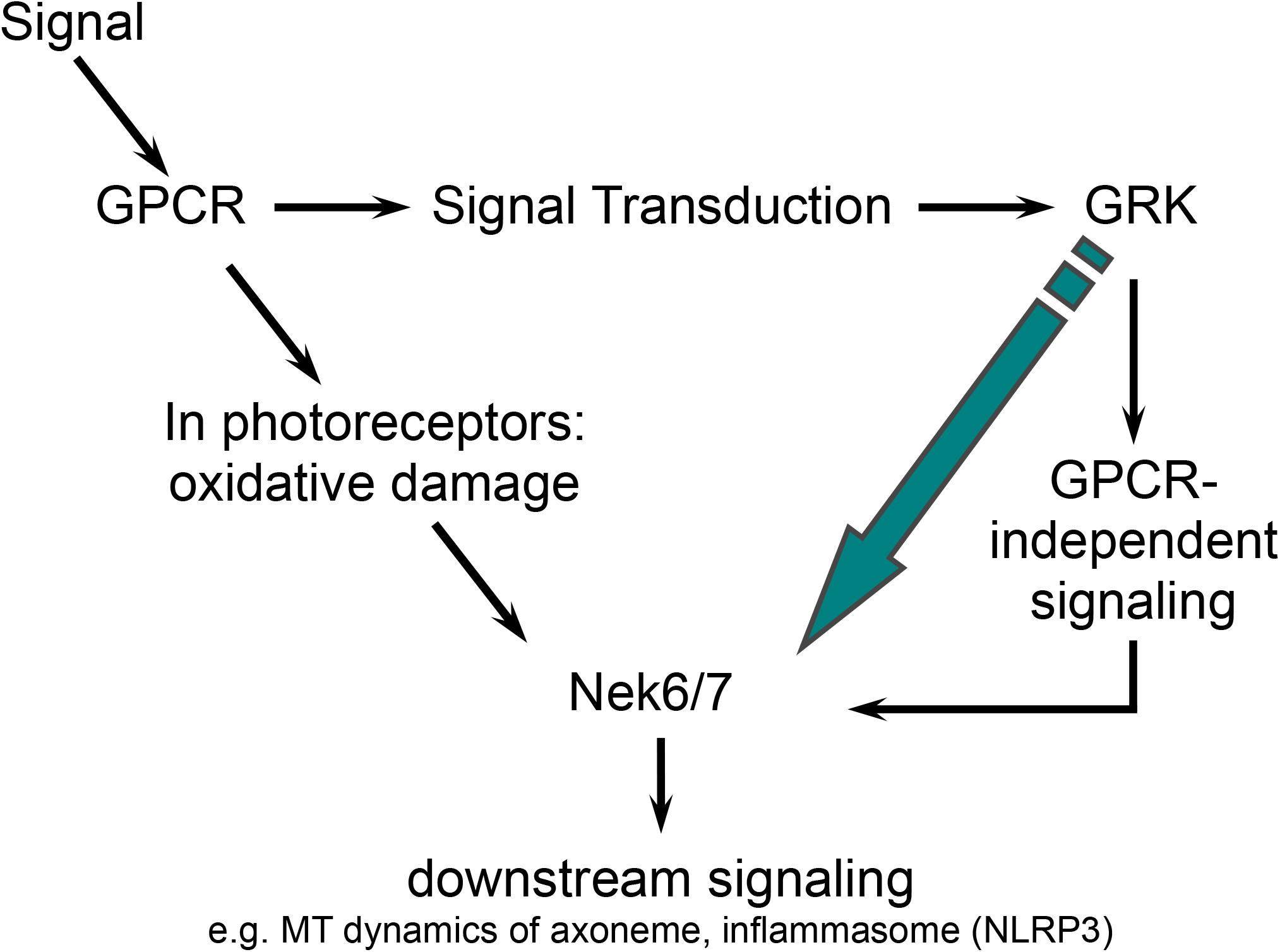
Proposed model for Nek6/7 kinase activation. GPCR, G-protein coupled receptor; GRK, G-protein coupled receptor kinase; Nek, NIMA-related kinase

## Materials and Methods

### Chemicals

NaCl, NaOH, Tris-HCl, methanol, isopropanol, phosphatase inhibitor cocktails I+II, protease inhibitor cocktail, 10x Tris-Glycine-SDS buffer, Tween 20, Triton X-100, glycerol, EDTA, EGTA, CHAPS, β-glycerolphosphate, NP-40, benzonase nuclease, urea, thiourea, D-amphetamine, cocaine, norepinephrine, buprenorphine, ampholyte high resolution pH 3–10, ampholyte high resolution pH 8–10, ampholyte high resolution pH 7–9 were purchased from Sigma-Aldrich, St. Louis. Phosphoric acid was purchased from Fisher, Pittsburgh. Nitrocellulose membranes, acrylamide/bis-acrylamide, TEMED, ammonium persulfate were from BioRad, Hercules. 1,4-dithiothreitol (DTT) and bovine serum albumin (BSA) for blocking were from Santa Cruz Biotechnology Inc, Santa Cruz. Immobilon Chemiluminescence HRP substrate and MilliQ Water (> 18.5 mΩ) were from Millipore, Temecula. VECTORSHIELD mounting media with DAPI was from Vector Laboratories, Burlingame. Dark- and light-adapted cow rod outer segments were from InVision BioResources, Seattle. HotStart-IT® SYBR® Green one-step qRT-PCR master mix kit was from usb, Cleveland. SuperScript® III First-Strand Sythesis SuperMix for qRT-PCR and primers were purchased from Invitrogen, Carlsbad.

### Antibodies

The phosphoNek6/7 antibody was a generous gift from Dr. Joan Roig Amorós (Cell Signaling Research Group, Institute for Research in Biomedicine, Barcelona, Spain). Nek6 antibody was from Abgent, San Diego. Nek7 antibody for IHC was purchased from Aviva Systems Biology, San Diego; Nek7 antibody for immunoblots was from Cell Signaling Technologies, Danvers. Centrin-1 antibodies were from Abcam, Cambridge. Rhodopsin (1D4) antibody, goat anti-rabbit IgG-HRP and goat anti-mouse IgG-HRP were from Santa Cruz Biotechnology Inc., Santa Cruz. Rabbit anti-beta actin antibody was from Bethyl Laboratories, Inc, Montgomery. Alexa Fluor® 488 goat anti-rabbit and Alexa Fluor® 555 goat anti-mouse IgG was from Invitrogen, Carlsbad.

### Mouse lines and husbandry

MF1 mice were purchased from Harlan, Somerville, NJ, and housed in a pathogen-free animal facility at Columbia University. Mice were used in accordance with the Statement for the Use of Animals in Ophthalmic and Vision Research of the Association for Research in Vision and Ophthalmology, as well as the Policy for the Use of Animals in Neuroscience Research of the Society for Neuroscience.

### Experimental procedure

MF1 and knockout mice were kept in a 12-hr light/dark cycle with food and water *ad libidum* until they reached experimental age (4-6 wks).

For light exposure experiments, MF1 and knockout mice were dark-adapted over night and sacrificed in complete darkness. Retinae were removed and placed in a Petri dish with 5 μl Ringer buffer to prevent drying-out. The Petri dish was placed under a light source and the retinae were exposed to light. The retinae were snap-frozen in liquid nitrogen and stored at −80 °C prior to use.

For drug experiments, MF1 mice were injected with drugs subcutaneously. Animals were sacrificed after three days with injections and their brains removed and frozen immediately on dry ice prior to use. Frozen brains were dissected on ice with a brain slicer with 1 mm spacing for striatum and substantia nigra enriched samples.

### qRT-PCR

Total RNA extraction was done using the RNeasy mini kit (Quiagen, Valencia, CA). The SuperScript® III First-Strand Sythesis SuperMix for qRT-PCR from Invitrogen was used according to manufacturer’s instructions to generate cDNA from total RNA extractions (Invitrogen, Carlsbad, CA). qRT-PCR was done using the HotStart-IT® SYBR® Green one-step qRT-PCR master mix kit manufacturer’s instructions and β-actin was used as internal control to normalize qRT-PCR results (Dheda et al., 2004). qRT-PCR was done with an Applied Biosystems 7500.

Settings for the PCR reactions:

1 cycle of: 50° C for 10 min
1 cycle of: 95° C for 2 min
45 cycles of: 94° C for 15 s, 60° C for 30 s, 72° C for 60 s, followed by melting curve analysis

The following primers were used for the PCR reaction:

*nek6* forward: AAGCAACTGAACCATCCGAATATCA
*nek6* reverse: CCTAGGGACCAGATGTCTGACTTGA
*nek7* forward: CCCTGAGAGAACCGTTTGGAAATAC
*nek7* reverse: CTGTCGTAGCTCCTCCGAATAGTGA
*nek8* forward: GCGTAACTGAGAAATGAGATGGAGAA
*nek8* reverse: CTTGCTCTTGCTGCTGAGGATCTTA
*nek9* forward: GGACGTCATCAAGAGTGGCTGTAGT
*nek9* reverse: ACATGATTATCCCCACAGGAGACCT
*β*-actin forward: GATGACCCAGATCATGTTTGAGACC
*β*-actin reverse: TAATCTCCTTCTGCATCCTGTCAGC

### 2D electrophoresis

Commercially available dark- and light-adapted cow rod outer segments were lysed with 2D lysis buffer (10 M urea, 4 M thiourea, 1 M DTT, 2% Triton X-100, 20% isopropanol, 10% glycerol, 1.6% ampholyte high resolution pH 8-10, 0.4% ampholyte high resolution pH 7-9, protease inhibitor cocktail, phosphatase inhibitor cocktail I+II). Lysates were treated with benzonase nuclease to remove DNA from the samples. Protein content of the samples was determined with the RC DC protein assay from BioRad. For the first dimension, samples were loaded onto isoelectric focusing gels with 75 μg total protein content. Gels were prefocused at 250 V for one hour and run at 450 V for 22 h. For the second dimension, gel rods were loaded onto 15% acrylamide gels. Proteins were transferred onto a 0.2 μm nitrocellulose membrane overnight at 4 V/cm. Membranes were blocked with 3% BSA in 150 mM NaCl, 100 mM Tris-HCl pH 7.5 and 0.5% Tween 20. Proteins were detected with the following antibodies against: Nek6, Nek7 and rhodopsin. Immune complexes were visualised using the Immobilon Western Chemiluminescence HRP substrate kit utilising goat-anti species-specific IgG-HRP antibodies and BioMax Light Film developed in a Konika SX-101A. The films were recorded with an AlphaImager with AlphaImager Software for Windows (AlphaInnotech, San Leandro, CA).

### Immunohistochemistry

Eyes of dark- and light-adapted mice were fixed in 4% paraformaldehyde for one hour at room temperature. For frozen sections: Eyes were incubated in sucrose in PBS (5% - 30%) and OCT in 30% sucrose (1:1) for one hour each before embedding in OCT. Frozen OCT blocks were cut in 5 μm sections and stored at −80° C prior to use. Sections of dark- and light-adapted eyes were placed on same slides. Sections were washed twice in PBS and PBST (PBS + 0.1% Triton X-100) for 15 min each before blocking with 5% normal serum solution for one hour at room temperature. Sections were incubated overnight at 4° C with the antibody solution. After washing in 5% normal serum solution, PBST and PBS twice for 5 min each, sections were incubated with Alexa Fluor® 488 and Alexa Fluor® 555 secondary antibodies for two hours at room temperature. Slides were mounted with VECTASHIELD mounting media with DAPI after washes in 5% normal serum, PBST and PBS for 5 min each at room temperature.

### Immunoblots

Tissue lysates were prepared with lysis buffer (20 mM Tris-HCl pH 7.4, 0.1% NP-40, 10% glycerol, 30 mM β-glycerolphosphate, protease inhibitor cocktail, phosphatase inhibitor cocktail I+II, 1 mM EDTA, 1 mM EGTA, 1 mM DTT). Protein concentration of the lysates was determined with Bradford assay. Lysates were separated with 50 μg total protein loading per lane on 15% acrylamide gels. Proteins were transferred onto a 0.2 μm nitrocellulose membrane overnight at 4 V/cm. Membranes were blocked with 3% BSA in 150 mM NaCl, 100 mM Tris-HCl pH 7.5 and 0.5% Tween 20. Proteins were detected with the following antibodies against: Nek6, Nek7, phosphoNek6/7. Immune complexes were visualised using the Immobilon Western Chemiluminescence HRP substrate kit utilising goat-anti species-specific IgG-HRP antibodies and BioMax Light Film developed in a Konika SX-101A. The bands were recorded quantitatively with densitometry at AlphaImager with AlphaImager Software for Windows (AlphaInnotech, San Leandro, CA).

### Experimental settings, statistics and software

All experiments were done in at least triplicates and repeated on different days. Results are shown as average ± stdev calculated with Microsoft Excel (Microsoft Corporation). Graphs were done in Microsoft Excel. 2D images were imaged and post-processed with AlphaImager Software for Windows (AlphaInnotech, San Leandro, CA). ImageJ/Fiji was used for post-processing immunohistochemistry images.

## Abbreviations

NIMA: never in mitose A
NEK: NIMA-related kinase
GPCR: G-protein coupled receptor
GRK: G-protein coupled receptor kinase
OPL: outer plexiform layer
IPL: inner plexiform layer
OS: outer segment
IS: inner segment
MT: microtubules

## Acknowledgements

The authors would like to thank Dr. R Allikmets and Dr. J Sparrow for their help and input with the manuscript. Thanks to Yael S Grossman for her help with the experiments. Thanks to Jean J Pak for her help with the post-processing of the IHC pictures. Thanks to Jason Chen and Frank Chen for providing the dark- and light-adapted retinae of the T−/−, RGS9−/− and RK−/− mice. We like to thank Dr. Joan Roig Amorós (Cell Signalling Research Group, Institute for Research in Biomedicine (IRB Barcelona) Barcelona, Spain) for the generous gift of the Nek6/7 phospho-antibody.

This work was supported by Research to Prevent Blindness, Burroughs-Wellcome Program in Biomedical Sciences, the Bernard Becker-Association of University Professors in Ophthalmology-Research to Prevent Blindness Award, Foundation Fighting Blindness, Dennis W. Jahnigen Award of the American Geriatrics Society, Joel Hoffman Fellowship, Gale and Richard Siegel Stem Cell Fund, Charles Culpeper Scholarship, Schneeweiss Stem Cell Fund, Irma T. Hirschl Charitable Trust, Barbara & Donald Jonas Family Fund, Eye Surgery Fund, and Bernard and Anne Spitzer Stem Cell Fund.

